# UCHIME2: improved chimera prediction for amplicon sequencing

**DOI:** 10.1101/074252

**Authors:** Robert C. Edgar

## Abstract

Amplicon sequencing generates chimeric reads which can cause spurious inferences of biological variation. I describe UCHIME2, an update of the popular UCHIME chimera detection algorithm with new modes optimized for high-resolution biological sequence reconstruction (“denoising”) and other applications. I show that chimera frequency correlates inversely with divergence, that error-free chimera prediction from sequence is impossible in principle, and that UCHIME2 achieves higher detection accuracy than previous methods.

---

Amplicon sequencing is widely used to survey biological sequence variation in applications including marker gene metagenomics^1^, immune system repertoire analysis^2^ and cancer genomics^3^. In such experiments, chimeric amplicons form when an incomplete DNA strand anneals to a different template and primes synthesis of a new template derived from two different biological sequences^4^. Recent chimera detection methods include ChimeraSlayer^4^, UCHIME^5^, DECIPHER^6^ and CATCh^7^. UCHIME classifies a query sequence by making a *model* from a concatenated pair of sub-sequences (segments) in a reference database. The query is predicted to be chimeric if the score of its alignment to the model exceeds a threshold. The reference database is provided by the user or constructed *de novo* from the reads. UCHIME2 modifies the original UCHIME algorithm by classifying a query as *unknown* if uncertainty is high due to missing reference data. Such queries would be classified, perhaps misleadingly, as “non-chimeric” by UCHIME. UCHIME2 also adds heuristics with parameter values optimized for different applications and introduces a new strategy designed for filtering denoised (error-corrected) amplicon sequences.

To investigate chimeras encountered in practice, I used Illumina reads of the currently popular V4 region of the 16S ribosomal RNA (16S) gene from high- and low-diversity microbial communities (soil and human vagina, respectively) and from two artificial communities (mock1 and mock2); see Supp. Note 1 for details. I extracted reads with < 1 expected errors^8^ and made OTUs at 97% identity using UCLUST^9^. Using its recommended 16S reference (the ChimeraSlayer Gold database), UCHIME predicted 9% of soil and 39% of vagina OTUs to be chimeric, while using SILVA^10^ predicted 57% and 53% respectively. The lower sensitivity of Gold vs. SILVA is explained by its smaller size (5.1k vs. 1.4M sequences), and the relatively better performance of Gold on vagina vs. soil is explained by the higher frequency of well-known species in vagina. The frequencies predicted using SILVA are probably underestimates and are surely better than using Gold (Supp. Notes 2 and 3), showing that at least half the soil and vagina OTUs are chimeric.

Earlier work^4–6^ characterized chimeras by parent divergence because detection is harder when parents are similar. However, a chimera can be arbitrarily close to one parent, e.g. if the other's segment is short. A better indication of detectability is the number of differences compared to the closest known non-chimeric sequence (*reference divergence*, *D*). Measured frequencies of chimeras with 1 ≤ *D* ≤ 20 are reported in Fig. 1, showing that frequency correlates inversely with *D* and a majority have *D* < 10. I also considered the identity of a segment with its closest reference sequence (*segment identity*, *S*). *S* is <100% when a parent is missing from the database, which reduces detectability.

**Fig. 1.**
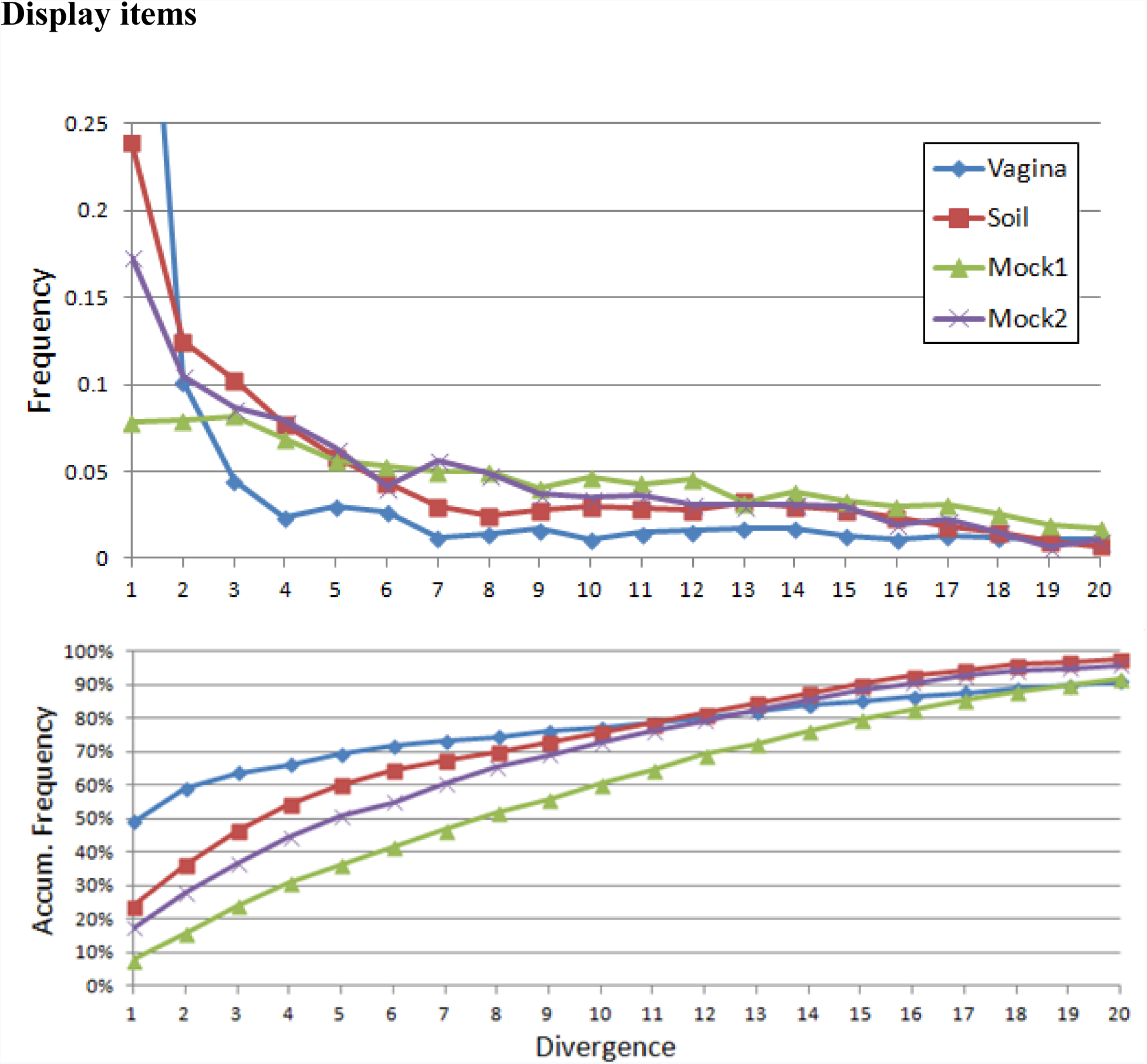
Chimera frequency as a function of divergence. Measured distributions for four communities with a wide range of diversities: soil (very high diversity), human vagina (low), and two mock communities (very low), which nevertheless exhibit similar distributions with an inverse correlation between divergence and frequency. See Supp Note 2 for methods. The horizontal axis is divergence, i.e. the number of differences between a chimera and the closest known non-chimeric sequence (which is almost certainly one of its parents in this analysis). The vertical axis is frequency (top) or accumulated frequency (bottom) calculated as a fraction of all chimeras found in a sample.. In all samples, a majority of chimeras have divergence < 10.

Previously published tests of reference-based chimera detection methods^4–6^ use simulations which assume, unrealistically, that parent sequences are known, i.e. *S* = 100%. To investigate the dependence of prediction accuracy on both *D* and *S*, I designed a new benchmark, CHSIMA, with simulated 16S and ITS chimeras with *D* = 1 to 10 and *S* = 90% to 100% (Supp. Note 5). False positives (FPs) were measured by dividing a chimera-free database into pairs (*splits*) with identities from 90% to 100%. I calculated sensitivity as the fraction of simulated chimeras that were correctly predicted and, unlike previously published tests, included false negatives (FNs) as well FPs in the error rate because both may be comparably harmful, especially when *D* > 3% where FNs cause spurious OTUs and FPs discard valid biological sequences. For assessments of DECIPHER and CATCh see Supp. Notes 4 and 6, which show DECIPHER to have very low sensitivity. CHSIMA results are given in Table 1 and Supp. Note 12. The *balanced* and *sensitive* modes of UCHIME2 have higher sensitivity than previous methods, with *balanced* having the lower overall error rate, and the *high-confidence* mode reports fewer false-positives.

**Table 1.** **CHSIMA test results on the 16S V4 region.** See Supp. Note 5 for details of the CHSIMA implementation, Note 11 for specifications of the methods and Note 12 for detailed results. *SegId* is the segment identity. Table entries are sensitivity (%) and total error rate (%) including both FPs and FNs. The highest sensitivity and lowest error rate for each *SegId* are underlined.

| Method | SegId=90%<br>*Sens (Err)* | SegId =95%<br>*Sens (Err)* | SegId =97%<br>*Sens (Err)* | SegId =99%<br>*Sens (Err)* | SegId =100%<br>*Sens (Err)* |
| --- | --- | --- | --- | --- | --- |
| U2-Denoised | 0.0 (50.0) | 0.0 (50.2) | 0.4 (52.0) | 6.6 (55.6) | <u>99.9</u> (<u>0.1</u>) |
| U2-Balanced | 47.3 (28.8) | 77.6 (<u>18.1</u>) | 87.3 (<u>10.8</u>) | 93.5 (<u>4.2</u>) | 98.0 (1.0) |
| U2-Sensitive | <u>76.6</u> (<u>23.4</u>) | <u>89.7</u> (26.2) | <u>93.6</u> (22.6) | <u>95.8</u> (12.2) | 99.4 (0.3) |
| U2-Specific | 12.7 (44.0) | 55.4 (23.4) | 75.1 (13.5) | 87.7 (6.3) | 95.5 (2.2) |
| U2-HighConf. | 0.7 (49.7) | 23.7 (38.5) | 52.7 (23.6) | 72.5 (13.8) | 86.2 (6.9) |
| UCHIME | 14.6 (43.2) | 58.5 (21.9) | 77.2 (12.8) | 88.3 (6.0) | 94.2 (2.9) |
| ChSlayer | 2.8 (48.6) | 29.7 (35.5) | 54.4 (23.6) | 68.9 (16.6) | 87.8 (6.5) |
| ChSlayer-kmer | 3.6 (48.2) | 32.1 (34.4) | 54.0 (23.5) | 65.7 (18.1) | 87.9 (6.5) |

The sensitivity of all tested methods falls rapidly with decreasing segment similarity. This is explained by chimeric models (*fakes*) of non-chimeric queries (Supp. Note 7), which are common and in the most challenging cases are exact matches (*perfect fakes*). Remarkably, I found that a large majority of non-chimeric sequences in the 99% identity splits have perfect fake models for the V4 region of 16S and the ITS1 and ITS2 fungal Internal Transcribed Spacer regions, and a third to a half in the 97% identity splits. Introducing heuristics to reduce FPs due to fakes therefore unavoidably causes an increase in FNs, especially for low-divergence chimeras. Error-free chimera/non-chimera classification from sequence is therefore impossible in principle, even in an ideal scenario where the reference database is complete and correct and the query sequence has no errors (Supp. Note 9). The problem of fakes is exacerbated when there are sequence errors or the database is incomplete, as is typically the case, noting that when *S* ≲ 100%, the probability of a non-chimera having a perfect fake model is very high (Supp. Note 7). These results, combined with the rapid increase in FNs with lower segment identities (Table 1), show that the choice of Gold as default for UCHIME was misguided; much better accuracy will usually be achieved with a large database such as SILVA due to the reduction in FNs.

## Methods

Given a query sequence *Q*, UCHIME2 uses the UCHIME algorithm to construct a model (*M*), then makes a multiple alignment of *Q* with the model and top hit (*T*, the most similar reference sequence). The following metrics are calculated from the alignment: number of differences *d_QT_* between *Q* and *T* and *d_QM_* between *Q* and *M*, the alignment score (*H*) using eq. 2 in Edgar *et al.* 2011. The fractional divergence with respect to the top hit is calculated as *divT* = (*d_QT_* – *d_QM_*)/|*Q*|. If *divT* is large, the model is a much better match than the top hit and the query is more likely to be chimeric, and conversely if *div_T_* is small, the model is more likely to be a fake. If *div_T_* > 3%, the chimera would cause a spurious OTU.

Deep sequencing enables resolution of amplicons with as few as one difference^11,12^. In marker gene applications, this can be accomplished by error-correction (*denoising*) using algorithms such as DADA2^12^ and UNOISE^8^ which predict the set of unique amplicon sequences from which the reads are derived. Many amplicons will be chimeric, and a post-processing step is therefore required to identify the non-chimeric subset. With this application in mind, I designed a *denoised de novo* (DDN) mode of UCHIME2 using a strategy similar to the *de novo* mode of UCHIME. Each sequence is compared with all amplicons having greater abundance on the assumption that parents are more abundant than chimeras because they undergo more rounds of amplification. If a perfect model is ^found (*d_QM_*=0*, d_QT_* > 0), the amplicon is classified as chimeric. This method is not heuristic in^ the sense that a perfect model will always be reported if one exists. Unlike UCHIME *de novo*, DDN will detect chimeras with divergences as low as a single difference. DDN compares an amplicon with all higher-abundance sequences, including those with chimeric models (which would be discarded by UCHIME), enabling detection of chimeras with three or more segments which form^13^ when a parent is itself chimeric. State-of-the-art denoisers achieve very low error rates^12^, giving amplicon sequences close to the ideal scenario described in the main text: error-free sequences and a complete reference database. Even in this scenario, false positives are possible because a valid biological variant with one difference has a perfect fake *de novo* model with a probability of a few percent (Supp. Note 9).

However, given that this probability is low and that *D*=1 chimeras are common (Fig. 1) I considered it preferable to classify any query with a perfect model as chimeric, allowing a few FPs due to perfect fakes rather than the many FN *D*=1 chimeras that would typically result from imposing more stringent criteria.

Accurate *de novo* chimera detection for high-resolution applications requires denoised amplicon sequences. Even if read quality filtering is applied, for example by requiring that the most probable number of errors implied by Phred scores is zero, many reads with one or more errors remain^8^. *De novo* chimera detection would then be severely degraded by the explosion in perfect fake models when *S* < 100% (Supp. Note 7).

For lower-resolution (OTU) clustering of quality-filtered (but not denoised) reads at 97% identity, UPARSE was shown^14^ to generate dramatically fewer chimeric OTUs on mock community data than pipelines which use UCHIME for chimera filtering. This was achieved despite the inherent limitations of chimera detection because UPARSE does not need to distinguish between low-divergence chimeras, reads with base call errors and biological variants with <3% differences—all are assigned to the closest OTU and their sequences *per se* are discarded. An OTU pipeline should therefore use UPARSE for clustering and chimera filtering unless reads are denoised, in which case using UCHIME2 DDN followed by OTU clustering with UCLUST is a reasonable alternative.

Results in the main text show that 100% segment similarity is needed for accurate detection with a user-supplied database, but this will rarely be possible in practice because reads often have errors and reference databases are incomplete and contain sequence errors and ambiguous bases. Noting that chimeras with *D*=1 are common, UCHIME2 classifies queries as *non-chimeric* only when there is a 100% identical reference sequence and otherwise as *chimeric* or *unknown*. I designed four different sets of parameters for reference-based chimeric classification, as follows. *Denoised* mode reports all chimeras when query sequences are error-free and the reference database is complete and correct. *Balanced* mode attempts to minimize both FPs and FNs, giving the lowest overall error rate. *Specific* mode prioritizes reducing FPs at the expense of increased FNs, with results similar to UCHIME. (Here, *unknown* is considered to be a FN if the query is a chimera). *High-confidence* mode further reduces FPs, at the cost of the highest overall error rate. *Sensitive* mode is designed to reduce FNs at the expensive of a higher FP rate. Parameter values for each mode are specified in Supp. Note 10. Noting that users rarely set non-default parameters even when command-line options are available, I did not set a default mode for UCHIME2, hoping that this would make users more aware of the limitations of reference-based chimera filtering by asking them to choose a compromise between sensitivity, false-positive and false-negative error rates that is most appropriate for addressing their biological questions.

## Author contributions

R.C.E. conceived of the study, performed the analysis and wrote the manuscript.

## UCHIME2: improved chimera prediction for amplicon sequencing

### Supplementary Notes

Note 1. Reads of soil, vagina and mock communities. Note 2. Measuring chimera divergences and frequencies.

Note 3. SILVA gives the best chimera frequency measurement. Note 4. Assessment of DECIPHER and CATCh.

Note 5. Design of the CHSIMA benchmark.

Note 6. Comments on the DECIPHER benchmark. Note 7. Fake and perfect fake models are common.

Note 8. Probability of a *de novo* perfect fake model.

Note 9. Error-free classification is impossible in principle. Note 10. Default parameters for UCHIME2 modes.

Note 11. Method names, software versions and command lines. Note 12. CHSIMA results.

Supplementary References

## Note 1. Reads of soil, vagina and mock communities

Reads for soil, human vagina and artificial community mock1 were taken from Kozich *et al.*^1^. The reads were downloaded from http://www.mothur.org/MiSeqDevelopmentData/, accessed 3rd Sept 2015.

Mock2 is MiSeq dataset 2 from Bokulich *et al*.^2^ The reads were kindly provided by Dirk Gevers at the Broad Institute. There is now a link to the data from the QIIME resource page (http://qiime.org/home_static/dataFiles.html, accessed 23rd June 2016).

## Note 2. Measuring chimera divergences and frequencies

In order to measure chimera divergence frequencies reliably, it is necessary to align sequences which can be classified as chimeric or not with high confidence against error-free parent sequences using a method that is unbiased with respect to divergence. This is challenging because it is not possible to identify sequencing error with 100% certainty, and error-free chimera detection is not possible, especially with noisy reads, and especially when divergence is low, as shown in the main text. This note describes a method which overcomes these challenges to make an accurate measurement of divergence frequencies.

### Reference database

For the mock1 and mock2 communities, I used the known 16S sequences for the species in the samples. For the soil and vagina samples, I used the 100 most abundant error-corrected amplicons generated by UNOISE^3^ as the reference database. The top 100 sequences are almost certainly correct (because denoising is most effective at high abundances) and a large majority of chimeras have parents in the top 100. Regardless, the subset of chimeras with top-100 parents is surely unbiased with respect to divergence frequencies.

### Query set

I searched for chimeras in reads after filtering at maximum one expected error^3^. I did not use error-correction because many chimeras are singletons, which are discarded by denoising methods including DADA2^4^ and UNOISE.

### Chimera detection algorithm

I used the denoised mode of UCHIME2 which is not heuristic in the sense that it is guaranteed to report all perfect models, i.e. all models with *d_QM_* = 0 and *d_QT_* > 0 (see Methods in main text) and will therefore identify all error-free chimeric sequences with parents in the reference database. Sequencing error in a chimeric read will almost certainly result in a false negative (because then *d_QM_* = (number of base call errors) > 0 except in pathological cases where errors reproduce a chimera by chance). The false-positive rate for error-free reads against error-free parents is estimated in Note 9 and found to be a few percent for *D*=1 and vanishingly small for *D* > 1.

### Despite known sources of error, divergence frequencies are accurately measured

This method has a number of well-understood sources of error. A measurement of divergence frequencies will be degraded only if these errors are biased with respect to divergence, otherwise they are benign. False negatives are caused by (1) unfiltered sequencing error and (2) parents missing from the reference database, neither of which is biased w.r.t. chimera divergence. False positives are rare and the causes should again have no bias w.r.t. divergence with the possible exception of *D*=1. Therefore, the method described in this note gives an accurate measurement of the chimera divergence distribution except that *D*=1 may be overestimated.

## Note 3. SILVA gives the best chimera frequency estimate

To investigate the coverage of the SILVA and Gold databases on the human vagina and soil samples, I measured the identities of the top hits of the 100 most abundant denoised amplicons (Fig. SN3.1). SILVA has ≥99% identity for 96% of the vagina amplicons and 75% of the soil amplicons, while Gold has ≥99% identity top hits for 50% and 6% of the amplicons, respectively. Referring to Table 1 in the main text, UCHIME2 in balanced mode has sensitivity 87% for identities of 97% or higher with an approximately equal number of FPs and FNs. Chimera frequencies from UCHIME2-balanced predictions with SILVA should therefore be superior to Gold, especially for soil, though will probably give an underestimate due to the lack of high identity reference sequences for ~25% of the high-abundance amplicons in soil.

**Fig. SN3.1.**
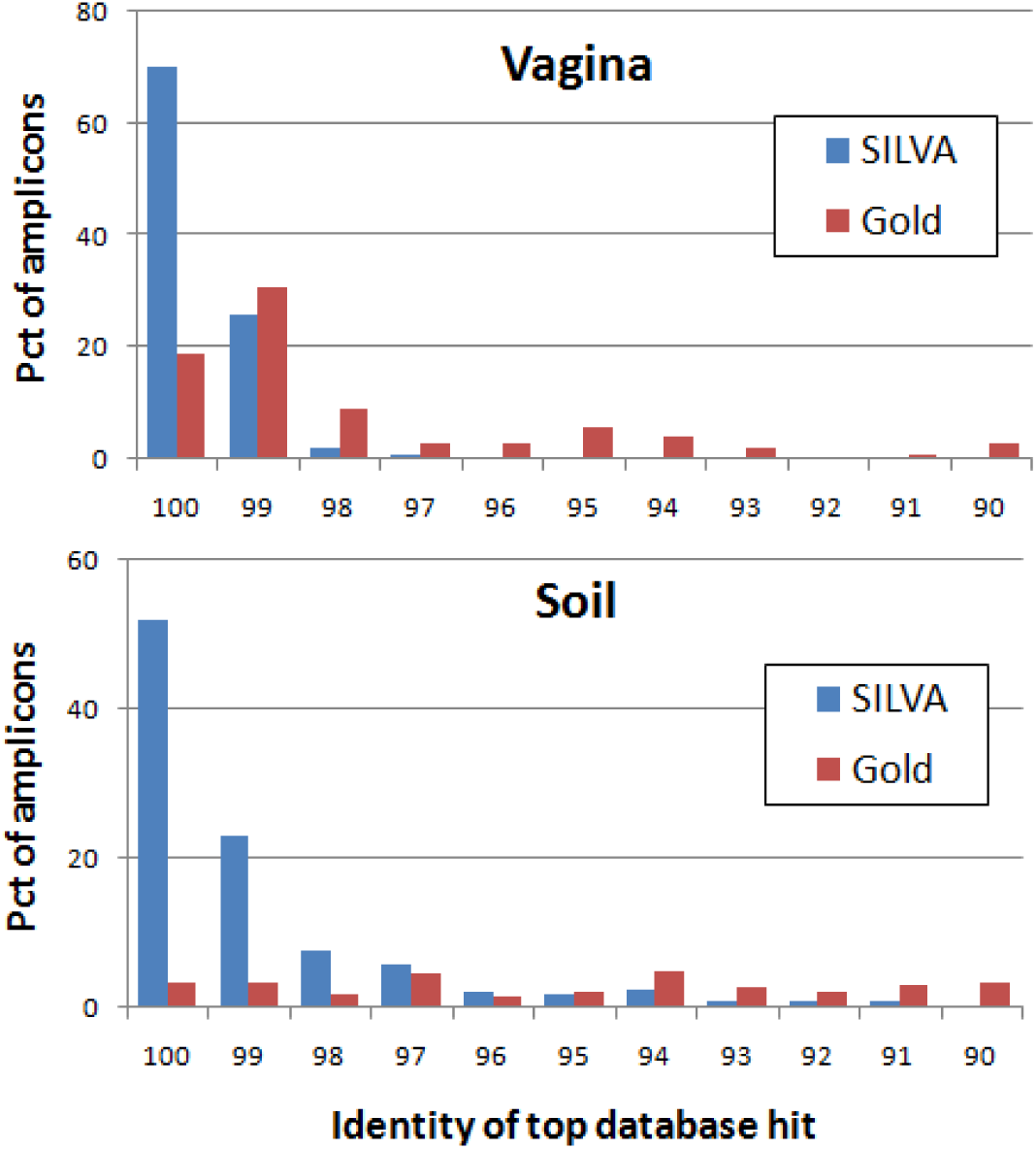
Top-hit identity distribution of the 100 most abundant denoised amplicons in soil and human vagina against the SILVA and Gold databases. SILVA top hits have ≥99% identity for 96% of the vagina amplicons and 75% of the soil amplicons, while Gold has ≥99% identity top hits for only 50% and 6%, respectively.

## Note 4. Assessment of DECIPHER and CATCh

CATCh^5^ is an ensemble classifier that generates a consensus result from predictions by UCHIME^6^, DECIPHER^7^ and other algorithms. I was unable to install the stand-alone version of DECIPHER or to configure DECIPHER to use a non-default reference database, which precluded running DECIPHER or CATCh on most benchmark tests including the ChimeraSlayer^8^ benchmark, the CHSIMA test described in the main text, or the SIM2 and CHSIM tests described in the UCHIME paper.

I tested DECIPHER and CATCh using the length-300 ChimeraSlayer test sequences (https://sourceforge.net/projects/microbiomeutil/files/ChimeraTestRegime/, retrieved 2nd Feb 2016). These include 2,500 length-300 two-segment chimeras (*Ch300*) constructed from parents in the Gold database and 4,770 length-300 segments (*Ctl300*) of Gold sequences, which are named isolates and thus known to be non-chimeric. I chose this test because it uses published benchmark data, the length is close to the currently popular V4 region (~250nt) and because it is reasonable to assume that all the Gold sequences are present in the reference databases used by DECIPHER and CATCh, noting that if parent sequences were not present, this would bias a test in favor of UCHIME and UCHIME2. For DECIPHER, I used its web server (http://decipher.cee.wisc.edu/FindChimeras.html, data submitted 1st Jul 2016). CATCh predictions were kindly provided by Mohamed Mysara.

In the ChimeraSlayer benchmark, divergence (*PDiv*) is measured between parents rather than the distance (*D*) between the chimera and its closest parent. *PDiv* provides an approximate upper bound *D* ≤ *PDiv*/2 because the closer parent is probably the one with the longer segment, which covers at least half of the chimera. Thus, *PDiv* ≤ 6% corresponds approximately to *D* ≤ 3%, the “harmful range" where chimeras will cause spurious OTUs in a typical analysis. Low-divergence chimeras are common (main text), so the “abundant range” *D* ≤ 5% or *PDiv* ≤ 10% is the most important in practice. Above *PDiv* = 10%, chimeras are relatively rare and are detected with high sensitivity by all tested methods except DECIPHER.

Results are shown in Table SN4.1 and in Figs. SN4.1 and SN4.2. The sensitivity of DECIPHER was found to be much lower than the other methods, while CATCh had sensitivity slightly higher than UCHIME. Both CATCh and DECIPHER reported several false positives despite that fact that all of the control sequences are segments of named isolate sequences, while UCHIME and all UCHIME2 modes reported no false positives as expected for known sequences. Note that this is *not* the same procedure used for the false positive tests described in the ChimeraSlayer and UCHIME papers which used a “leave-one-out" strategy (i.e., the query sequence was omitted from the database). In this case, I could not change the databases for DECIPHER or CATCh so I tested all methods using “leave-all-in”, i.e. without deleting the query sequence, as in the CHSIMA false positive tests with *SegId*=100% (Note 5).

**Fig SN4.1.**
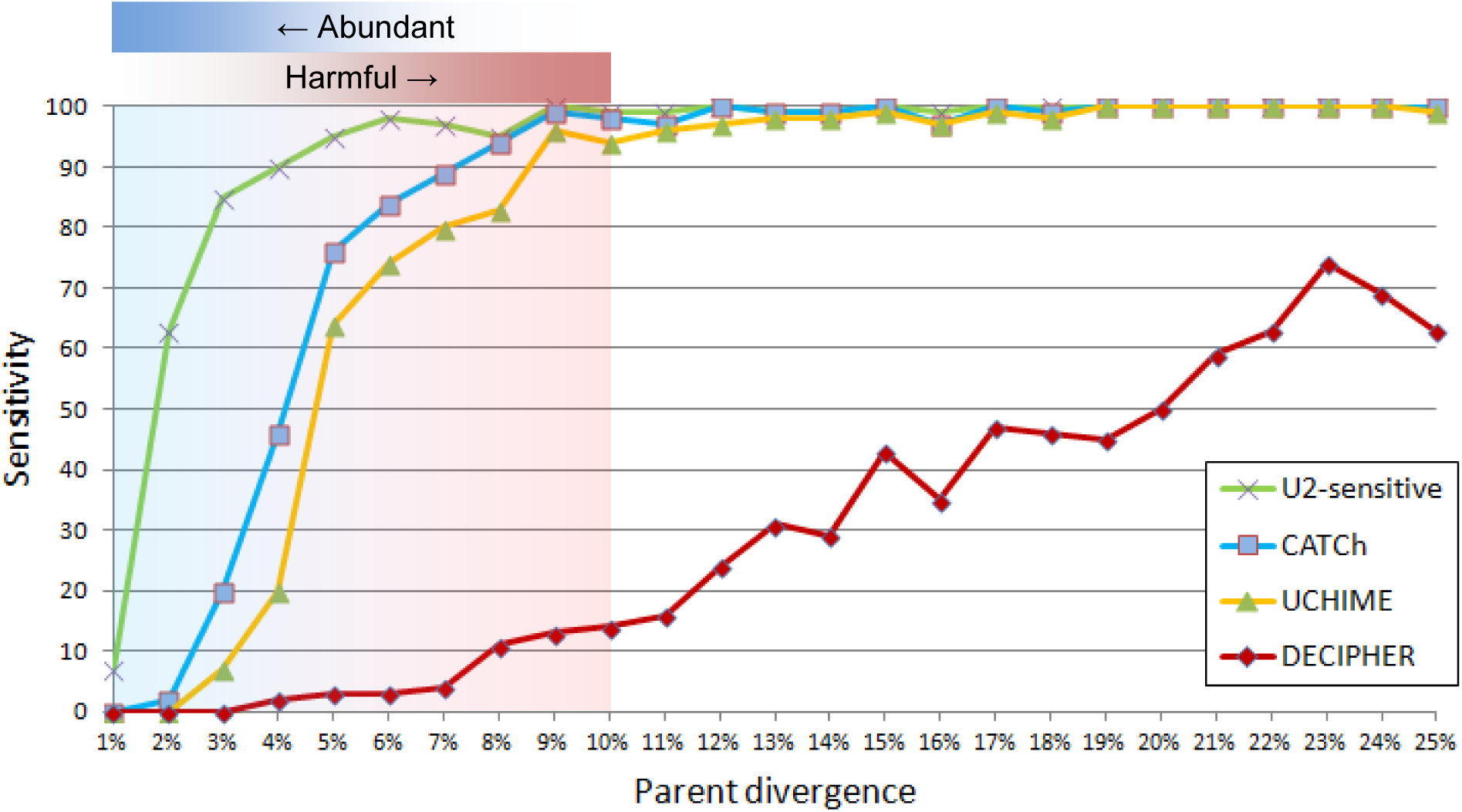
Sensitivity on the Ch300 set. The chart shows sensitivity of UCHIME2 in sensitive mode (U2-sensitive), CATCh, UCHIME and DECIPHER as a function of parent divergence (*PDiv*).

**Fig. SN4.2.**
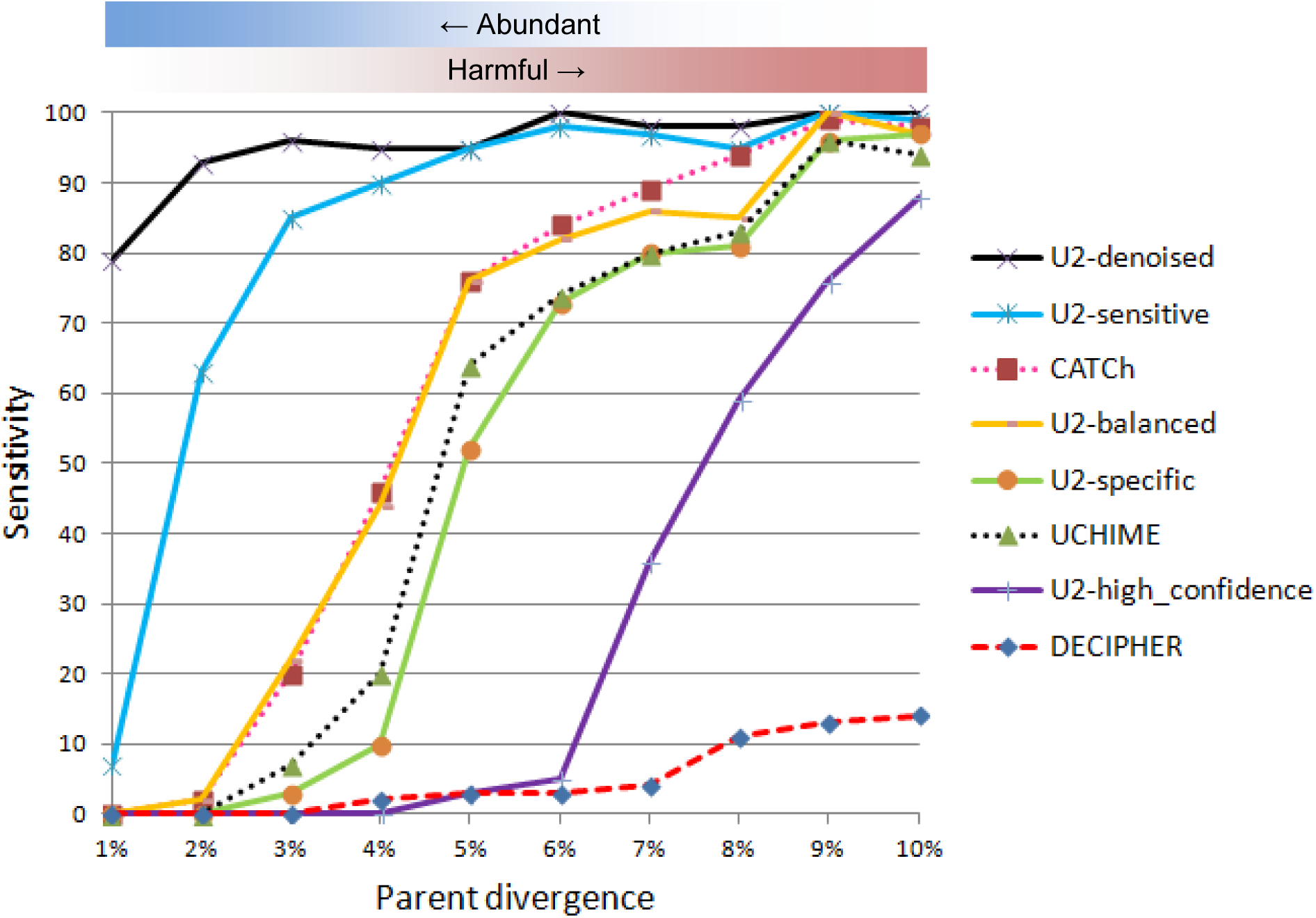
Sensitivity on low-divergence chimeras in Ch300. Results showing only chimeras with *PDiv* ≤ 10% as these are the most abundant, hardest to detect, and most harmful in a typical analysis. U2-*m* is UCHIME2 in mode *m*. UCHIME2 in denoised mode is guaranteed to find all perfect chimeras; its sensitivity is nevertheless <100% because 38 chimeras in the Ch300 set are 100% identical to reference sequences.

**Table SN4.1.** **False positives on the Ctl300 test.** The table shows the number of false positives reported by each method on 4,770 length-300 segments of high-quality named isolate sequences. UCHIME and UCHIME2 report no false positives because all of the segments have 100% identical full-length matches in the reference database.

| Method | FPs | FP (Pct) |
| --- | --- | --- |
| CATCh | 154 | 3.2% |
| DECIPHER | 43 | 0.9% |
| UCHIME | 0 | 0% |
| UCHIME2 (all modes) | 0 | 0% |

UCHIME2 in denoised mode is not heuristic in the sense that it will report all query sequences not found in the reference database for which a perfect chimeric model can be constructed. Its sensitivity is <100% on Ch300 (Fig. SN4.2) because 38 of the simulated chimeras have hits to Gold with 100% identity, e.g. L300chmraD1_S000440923_5642-6199:6200-6518_S000382508 matches S000842173, illustrating that error-free prediction is impossible in principle (Note 8).

## Note 5. Design of the CHSIMA benchmark

My goal with CHSIMA was to implement a benchmark for reference-based methods that measures accuracy with varying chimera divergence (*D*) and varying segment similarity (*S*). Previous benchmarks tested variation with *D* but made the unrealistic assumption that parent sequences are always known, i.e. *S*=100%.

For a given region (e.g., V4 or ITS1), I extracted the region from the reference database, ^giving sequence set *R*. For each *S*, I constructed two subsets (*splits X_S_* and *Y_S_*) of *R* with^ approximately equal numbers of sequences such that each sequence in one split had a top ^hit with identity *S* in the other split (Fig. SN5.1). In the special case *S*=100%, *X*^100 ^= *Y*^100 ^= a^ random subset of 1,000 sequences from *R*.

**Fig. SN5.1.**
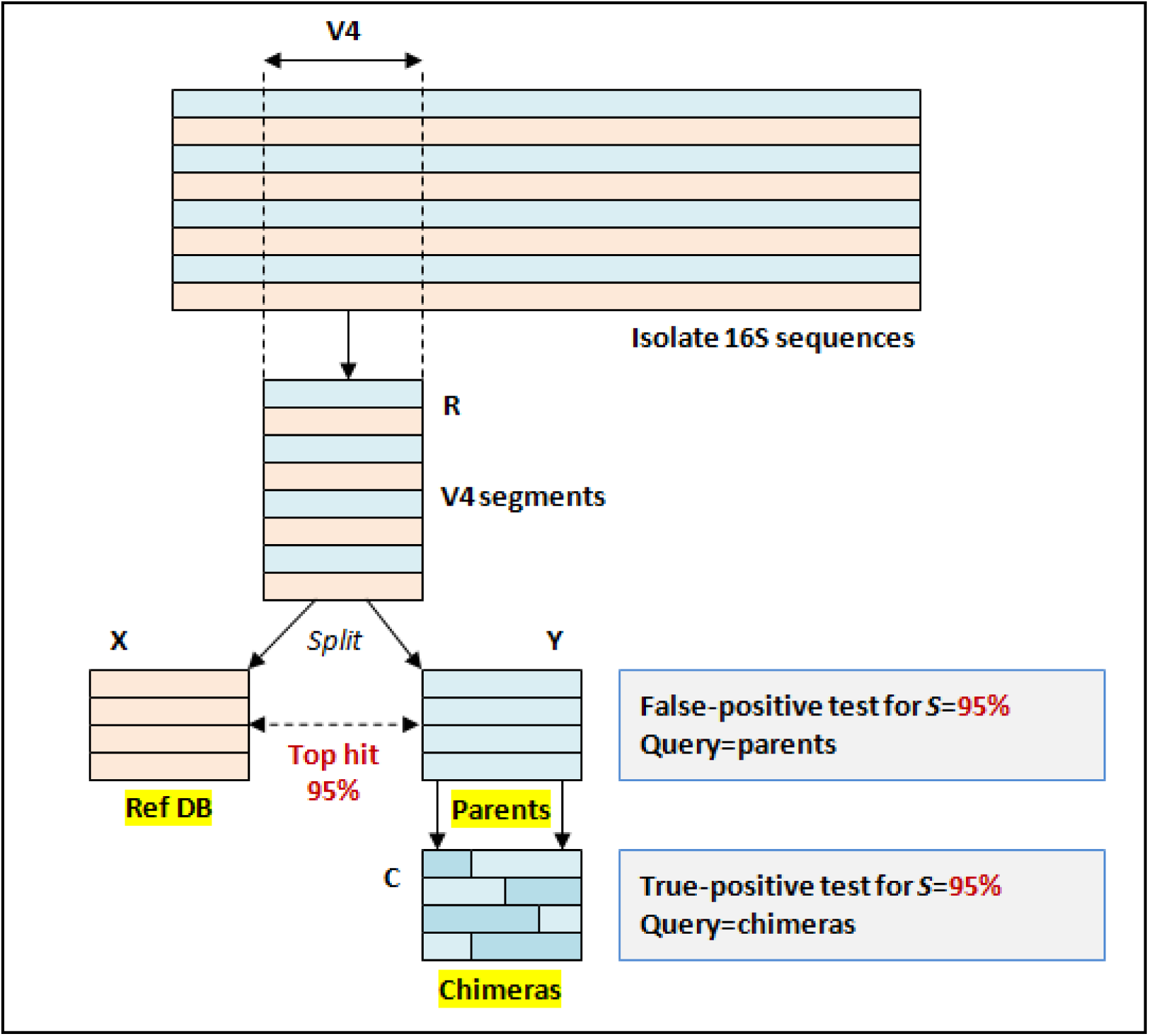
Construction of the CHSIMA benchmark. The figure shows the method used to construct a typical subset of CHSIMA using the V4 region and *S*=95% as an example.

For each pair of splits, I constructed a set of two-segment simulated chimeras *C_S_* from parent sequences in *Y_S_*, generating 100 chimeras for each *D* = 1, 2 … 10 for a total of 1,000.

True positives (TPs) are measured with *X_S_* as the reference database and *C_S_* as the query set. Sensitivity is the fraction of chimeras that are successfully detected and the false-negative (FN) rate is the fraction not detected.

False positives (FPs) are measured with a random subset of 1,000 sequences from *Y_S_* as the query and *X_S_* as the database. This measures the FP rate with a known distance (*S*) between the database and the query sequences, in contrast to the leave-one-out method used in the ChimeraSlayer and UCHIME papers where the distance has varying values <100% but is not otherwise characterized.

I defined sensitivity and error rates for a given *S* as follows:

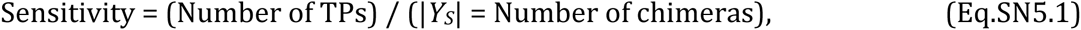

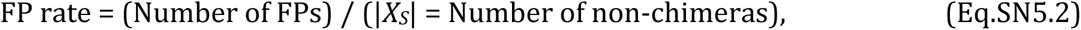

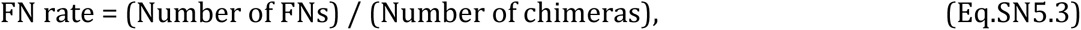

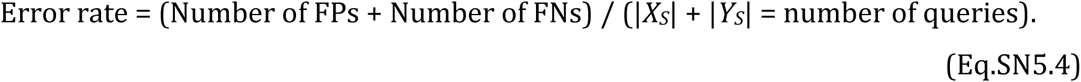

In most cases, |*X_S_*| = |*Y_S_*| = 1,000, in which case the overall error rate is the mean of the FP rate and FN rate:

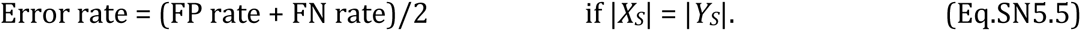

Sensitivity and error rates may be given for each *D* or averaged over all *D* = 1, 2 … 10.

Previous benchmarks^5–8^ did not include false negatives in reported error rates. While the false negative rate is readily calculated as FN rate = (100% – Sensitivity), I felt it was more informative to include false negatives in reported total error rates because of the atypical importance of false-negative chimera predictions. A sensitivity of 90% and false-positive error rate of 2% would sound pretty good for a typical informatics algorithm, but here if you have 1,000 OTUs then you will end up with 10% = 100 spurious OTUs due to false-negative chimeras in addition to the 20 good OTUs lost due to false positives. In my opinion, it is more informative to express this as a 12% error rate so that a casual reader or user does not overlook the importance of FNs. For another example, the sensitivity of DECIPHER for 300nt chimeras with *S*=100% and *PDiv* ≤ 10 is 5% (Fig. SN4.2) with a FP rate of 1% (Table SN4.1) so the large majority of harmful chimeras will be missed by DECIPHER. In my opinion, quoting the overall error rate per query (95% + 1%)/2 = 48% gives a better indication of the poor performance of the algorithm on this data.

## Note 6. Comments on the DECIPHER benchmark test

I was not able to reproduce results in the DECIPHER paper because some details are not specified and I was unable to install the stand-alone version of DECIPHER.

The DECIPHER paper does not fully explain how false positives were measured, in particular which reference database was used for each algorithm. The authors state “[t]he 2,000 parent sequences in the data set were … evaluated to estimate the rate of false-positive detections … with fs_DECIPHER having the lowest false positive rate (0.15%), followed by ChimeraSlayer (0.70%), WigeoN (0.85%), Uchime (1.5%), and ss_DECIPHER (1.6%)." UCHIME has a zero false positive rate for query sequences having a 100% identical full-length match, so the database used for UCHIME was missing at least 1.5% = 30 of the full-length sequences for the 2,000 parents; in fact, probably many more, given its low FP rate. The Gold database does not contain all of the parents, and if Gold was used with UCHIME this could explain why its FP rate is >0 and raises the question of whether DECIPHER was run with its own default database which does include all the parent sequences and would bias the results in favor of DECIPHER.

Another issue with the DECIPHER benchmark arises from reference sequences with missing bases at the 5' and 3' terminals. While the RDP isolate and Gold sequences are sometimes considered to be “full-length", in fact they often do not cover the complete gene (see Fig. SN6.1 for an example). With default settings, UCHIME allows terminal gaps in query sequences but not reference sequences (see discussion of Global-X alignments and Fig. 1 in Edgar *et al.* 2011). This may partly explain the anomalously low sensitivity of UCHIME on the DECIPHER benchmark test.

**Fig. SN6.1.**
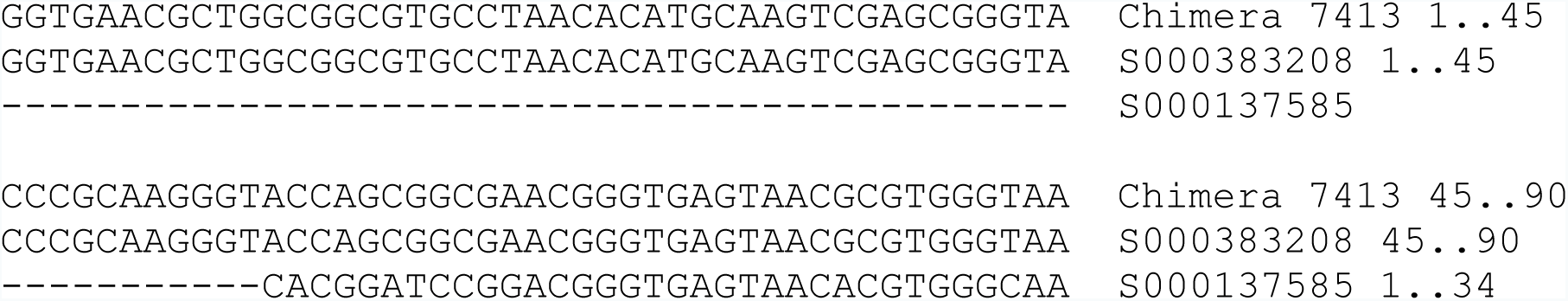
Missing reference sequence bases. Chimera 7413 in the TwoPart-1-Chimeras.txt subset of the DECIPHER benchmark is constructed from two segments of length 639 and 104 which are substrings of parent sequences S000383208 and S000137585 in the Gold database. The alignment has terminal gaps in S000137585 due to the incomplete sequence of S000137585, and is therefore rejected by UCHIME with default settings (see Edgar *et al.* 2011, Fig. 1 and discussion of Global-X alignments). I interpret this apparent false negative as an artifact of the benchmark implementation rather than a defect of the algorithm *per se* because in practice query sequences are usually next-generation reads which are fully covered by reference sequences. Applications where terminal gaps in a reference sequence might be expected, such as screening a reference database for chimeras, are much less common and are supported by UCHIME by setting appropriate non-default options.

## Note 7. Fake and perfect fake models are common

Given a query sequence *Q*, a *chimeric model* is a pair of reference sequence segments (*A*, *B*) concatenated together that have divergence > 0, i.e. are more similar to *Q* than the most similar sequence in the database (the *top hit*, *T*). If *Q* is not chimeric, the model is *fake*. A *perfect fake model* is a fake model that is identical to *Q*.

To give an indication of the frequency of fake models found by UCHIME2 I counted the number of alignments with a positive score, ignoring all other thresholds. Results are shown in the “Fakes” column in Table SN7.1. These are underestimates of the number of fake models that could be constructed, noting for example that this method fails to find many perfect fake models with *S*=99% found by UCHIME2-denoised (Table SN7.1).

All queries with at least one perfect fake model are identified by UCHIME2 in denoised mode because the algorithm is not heuristic, in the following sense: if one or more chimeric models exist, and the query sequence is not present in the database, then a model will be reported. Results for CHSIMA are shown in Table SN7.2 (see Note 5 for explanation of CHSIMA). This table can be interpreted as giving the false-positive error rate of UCHIME2-denoised, or equivalently as giving the frequency of perfect fake models for each segment identity (*SegId*, *S*). For all regions except full-length 16S, a large majority of sequences have a perfect fake model with *S*=99% and between a third and a half with *S*=97%. At *S*=99%, 10% of full-length 16S sequences have a perfect fake model.

The observation that both fakes and perfect fakes are very common implies that chimeras cannot be reliably distinguished from non-chimeras by any conceivable reference-based algorithm. In the *de novo* case, sequence abundances provide additional evidence which is predictive but not definitive. Similarly, it is not possible to screen large reference databases such as SILVA^9^ and Greengenes^10^ for chimeras with high accuracy. Low-divergence chimeras are common, and if the parents of a low-divergence chimera are present in the database, an algorithm such as UCHIME2-denoised can discover the model, but would not be able to reliably determine whether the model was fake because there is no evidence either way in the sequence. If its parents are not present, then the sensitivity and/or error rate of all algorithms necessarily increase rapidly with decreasing *S* and again, if a model is found, a chimera cannot be reliably distinguished from a correct biological sequence with a perfect or imperfect fake model.

**Table SN7.1.** **Fake and perfect fake models are common.** Number of fake models found by UCHIME-sensitive and number of perfect fake models found by UCHIME2-denoised. For all regions except full-length 16S, a large majority of sequences have a perfect fake model with *S*=99% and between a third and a half with *S*=97%.

| Region | SegId | Nr seqs in $X_S$ | Fakes | Perfect Fakes |
| --- | --- | --- | --- | --- |
| **V4**<br>**(~250nt)** | 99% | 1000 | 640 (64%) | 972 (97%) |
|  | 97% | 1000 | 775 (78%) | 483 (48%) |
|  | 95% | 1000 | 830 (83%) | 78 (8%) |
|  | 90% | 462 | 419 (91%) | 0 |
| **V3-V5**<br>**(~550nt)** | 99% | 1000 | 620 (62%) | 613 (61%) |
|  | 97% | 1000 | 755 (76%) | 374 (37%) |
|  | 95% | 1000 | 851 (85%) | 0 |
|  | 90% | 483 | 432 (89%) | 0 |
| **16S (full)** | 99% | 1000 | 642 (64%) | 99 (10%) |
|  | 97% | 1000 | 783 (78%) | 0 |
|  | 95% | 1000 | 888 (89%) | 0 |
|  | 90% | 655 | 607 93%) | 0 |
| **ITS1** | 99% | 1000 | 201 (20%) | 679 (68%) |
|  | 97% | 1000 | 366 (37%) | 340 (34%) |
|  | 95% | 1000 | 553 (55%) | 303 (30%) |
|  | 90% | 890 | 580 (65%) | 7 (8%) |
| **ITS2** | 99% | 1000 | 177 (18%) | 761 (76%) |
|  | 97% | 1000 | 413 (41%) | 504 (50%) |
|  | 95% | 1000 | 594 (59%) | 237 (24%) |
|  | 90% | 969 | 726 (75%) | 85 (9%) |
| **ITS (full)** | 99% | 1000 | 302 (30%) | 688 (69%) |
|  | 97% | 1000 | 594 (59%) | 496 (50%) |
|  | 95% | 1000 | 713 (71%) | 0 |
|  | 90% | 1000 | 808 (81%) | 0 |

## Note 8. Probability of a *de novo* perfect fake model

A *de novo* detection algorithm constructs a reference database from sequences found in the reads. The observation that perfect fake models are common (Note 7) raises the question of how often true biological variants with a few substitutions could cause false positives due to perfect fake models in a *de novo* database. To investigate this, I extracted the top 100 denoised amplicons for the soil and vagina samples and generated 100,000 random variants of these amplicons with 1, 2 … 5 substitutions. I then used UCHIME2-denoised with the variants as a query and the top 100 denoised amplicons as a reference to determine the fraction of variants which had a perfect chimeric model. This simulates the problem faced by the post-processing step in a denoising pipeline which attempts to predict which of the error-correct amplicon sequences are chimeric.

Results are shown in Table SN8.1, which show that a biological variant with one difference has a probability of a few percent of having a perfect fake model while a variant with two or more differences is very unlikely to have a perfect fake model. This shows that a few percent of low-abundance biological variants with a single substitution will be discarded as false positive chimeras. Biological variants with two or more substitutions are very unlikely to have perfect fake models.

**Table SN8.1.** **Probability that a biological variant has a perfect fake model.** Columns show the measured frequency of random variants with a given number of substitutions that have a perfect fake model. This shows that a biological variant with one difference has a probability of a few percent of having a perfect fake model.

| Sample | 1 sub | 2 subs | 3 subs | 4 subs |
| --- | --- | --- | --- | --- |
| **Soil** | 2.5% | 0.02% | 0 | 0 |
| **Vagina** | 5.5% | 0.1% | 0 | 0 |

## Note 9. Error-free chimera detection is impossible in principle

As discussed in Note 7, a *perfect fake model* is a pair of reference sequence segments (*A*, *B*) concatenated together that reproduce a query sequence *Q* which is known to be a correct biological sequence. If the reference database contains *A* and *B* but not *Q*, then it is impossible in principle for a reference-based algorithm to distinguish a chimeric amplicon *C*_*AB*_ = *A*+*B* from an amplicon derived from the correct biological sequence *Q* because the sequences of *C_AB_* and *Q* are identical. Perfect fake models are surprisingly common (Note 7).

A sequence which is found in the reference database and also has a perfect chimeric model constructed from two other reference sequences cannot be reliably classified for essentially the same reason -- the sequences of a potential chimera and a correct biological sequence are identical. The results of Note 7 show that such cases are very common, especially for the currently popular regions V4 and ITS. Consider *R_99_* = *X*_99_+ *Y*_99_for the V4 region. 97% of the sequences in *X*_99_have a perfect fake model in *Y*_99_, which implies that conversely ~97% of the sequences in *Y*_99_have perfect fake models in *X*_99._Combining the two splits, it follows that at least ~97% of the sequences in *R*_99_have perfect fake models constructed from other sequences in *R*_99_. The problem of perfect fake models is thus not solved in the ideal case where all biological sequences are known, noting that a set of denoised amplicons is a good approximation to this scenario.

In the *de novo* case, the unique sequence abundances of *A*, *B* and *Q* provide additional evidence which is predictive but not definitive because low-abundance variants with perfect chimeric models may be due to correct biological sequences with fake models (Note 8), to chimeric amplicons, or reads with incorrect bases due to uncorrected sequencing error, or polymerase copying mistakes. Since perfect fake models are relatively rare in the *de novo* case, especially when there multiple substitutions, UCHIME2-denoised-*de-novo* (DDN) classifies a sequence as chimeric if it has a perfect model and the parent sequences are more abundant.

## Note 10. Default parameters for UCHIME2 modes

**Table SN10.1.** Default parameters.

|  | *minh* | *mindiffs* | *mindivt* | *xa* | *xn* | *mindqt* | *maxdqm* |
| --- | --- | --- | --- | --- | --- | --- | --- |
| **denoised** | *n/a* | *n/a* | *n/a* | *n/a* | *n/a* | 1 | 0 |
| **high_confidence** | 0.80 | 3 | 1.0% | 1 | 8 | *n/a* | *n/a* |
| **specific** | 0.35 | 3 | 1.0% | 1 | 8 | *n/a* | *n/a* |
| **balanced** | 0.30 | 3 | 1.0% | 1 | 5 | *n/a* | *n/a* |
| **sensitive** | 0.15 | 2 | 0.5% | 1 | 4 | *n/a* | *n/a* |

## Note 11. Method names, versions and command lines

Source code and Linux binary for UCHIME2 is provided in the supplementary files. Command line options are given in the README.TXT file.

ChSlayer is the mothur^11^ implementation of ChimeraSlayer with default parameters. I used v1.36.1 of mothur. Typical commands:

~~~
mothur “#align.seqs(candidate=q.fasta, template=gold.align, processors=6)”
mothur “#chimera.slayer(fasta=q.align, template=gold.align, processors=6)”
~~~

ChSlayer-kmer is the same as ChSlayer plus the search=kmer option.

DECIPHER was run using its web server http://decipher.cee.wisc.edu/FindChimeras.html selecting the “short-length sequences" option for sequences < 1,000nt, “full-length sequences" for sequences ≥ 1,000nt. Data was submitted 1st Jul 2016.

CATCh results were kindly provided by M. Mysara.

## Note 12. CHSIMA results

**Table SN12.1.** **Summary of CHSIMA results for each region.** Sensitivity, FP and FN rates are given as percentages.

| ===== 16S (full) ===== |  |  |  |  |  |  |  |  |  |  |  |  |  |  |  |
| --- | --- | --- | --- | --- | --- | --- | --- | --- | --- | --- | --- | --- | --- | --- | --- |
| Algo | Sens | S=90<br>FP | FN | Sens | S=95<br>FP | FN | Sens | S=97<br>FP | FN | Sens | S=99<br>FP | FN | Sens | S=100<br>FP | FN |
| U2-Denoised | 0.0 | 0.0 | 100.0 | 0.0 | 0.0 | 100.0 | 0.0 | 0.0 | 100.0 | 0.0 | 9.9 | 100.0 | 94.3 | 0.0 | 5.7 |
| U2-Sensitive | 4.4 | 0.6 | 95.6 | 36.1 | 2.0 | 63.9 | 64.0 | 1.4 | 36.0 | 83.4 | 0.1 | 16.6 | 98.2 | 0.0 | 1.8 |
| U2-Balanced | 29.5 | 4.6 | 70.5 | 72.8 | 7.9 | 27.2 | 84.1 | 5.5 | 15.9 | 94.1 | 0.2 | 5.9 | 98.9 | 0.0 | 1.1 |
| U2-Specific | 6.6 | 0.7 | 93.4 | 51.3 | 2.7 | 48.7 | 73.8 | 2.1 | 26.2 | 90.3 | 0.2 | 9.7 | 98.8 | 0.0 | 1.2 |
| U2-HiConf | 0.4 | 0.1 | 99.6 | 14.1 | 0.7 | 85.9 | 43.2 | 0.8 | 56.8 | 70.3 | 0.0 | 29.7 | 98.3 | 0.0 | 1.7 |
| UCHIME | 14.3 | 0.4 | 85.7 | 66.5 | 1.7 | 33.5 | 84.3 | 3.3 | 15.7 | 92.9 | 0.3 | 7.1 | 96.0 | 0.0 | 4.0 |
| ChSlayer | 17.7 | 0.4 | 82.3 | 67.3 | 2.5 | 32.7 | 81.4 | 2.1 | 18.6 | 88.6 | 0.6 | 11.4 | 92.5 | 0.6 | 7.5 |
| ChSlayer-kmer | 23.8 | 0.4 | 76.2 | 70.3 | 2.6 | 29.7 | 83.0 | 2.7 | 17.0 | 90.1 | 0.6 | 9.9 | 93.4 | 0.8 | 6.6 |
| ===== V3-V5 ===== |  |  |  |  |  |  |  |  |  |  |  |  |  |  |  |
| Algo | Sens | S=90<br>FP | FN | Sens | S=95<br>FP | FN | Sens | S=97<br>FP | FN | Sens | S=99<br>FP | FN | Sens | S=100<br>FP | FN |
| U2-Denoised | 0.0 | 0.0 | 100.0 | 0.0 | 0.0 | 100.0 | 11.5 | 37.4 | 88.5 | 38.8 | 61.3 | 61.2 | 100.0 | 0.0 | 0.0 |
| U2-Sensitive | 11.2 | 0.1 | 88.8 | 63.6 | 1.9 | 36.4 | 82.4 | 5.0 | 17.6 | 92.9 | 1.2 | 7.1 | 98.5 | 0.0 | 1.5 |
| U2-Balanced | 38.1 | 1.0 | 61.9 | 76.4 | 10.7 | 23.6 | 89.1 | 10.9 | 10.9 | 94.8 | 2.3 | 5.2 | 98.6 | 0.0 | 1.4 |
| U2-Specific | 11.2 | 0.1 | 88.8 | 63.6 | 1.9 | 36.4 | 82.4 | 5.0 | 17.6 | 92.9 | 1.2 | 7.1 | 98.5 | 0.0 | 1.5 |
| U2-HiConf | 2.2 | 0.0 | 97.8 | 37.7 | 0.7 | 62.3 | 65.9 | 1.8 | 34.1 | 86.9 | 0.5 | 13.1 | 97.9 | 0.0 | 2.1 |
| UCHIME | 15.7 | 0.2 | 84.3 | 68.6 | 4.1 | 31.4 | 84.5 | 13.3 | 15.5 | 92.3 | 5.2 | 7.7 | 95.8 | 0.0 | 4.2 |
| ChSlayer | 13.9 | 0.4 | 86.1 | 62.6 | 3.2 | 37.4 | 77.1 | 6.5 | 22.9 | 84.8 | 3.9 | 15.2 | 92.5 | 2.5 | 7.5 |
| ChSlayer-kmer | 20.6 | 1.4 | 79.4 | 65.6 | 3.0 | 34.4 | 65.8 | 4.0 | 34.2 | 86.6 | 5.2 | 13.4 | 93.0 | 3.0 | 7.0 |
| ===== V4 ===== |  |  |  |  |  |  |  |  |  |  |  |  |  |  |  |
| Algo | Sens | S=90<br>FP | FN | Sens | S=95<br>FP | FN | Sens | S=97<br>FP | FN | Sens | S=99<br>FP | FN | Sens | S=100<br>FP | FN |
| U2-Denoised | 0.0 | 0.0 | 100.0 | 0.0 | 0.5 | 100.0 | 0.4 | 4.5 | 99.6 | 6.6 | 17.8 | 93.4 | 99.9 | 0.0 | 0.1 |
| U2-Sensitive | 76.6 | 23.4 | 23.4 | 89.7 | 42.1 | 10.3 | 93.6 | 38.8 | 6.4 | 95.8 | 20.1 | 4.2 | 99.4 | 0.0 | 0.6 |
| U2-Balanced | 47.3 | 4.9 | 52.7 | 77.6 | 13.8 | 22.4 | 87.4 | 8.9 | 12.6 | 93.5 | 2.1 | 6.5 | 98.0 | 0.0 | 2.0 |
| U2-Specific | 12.7 | 0.6 | 87.3 | 55.4 | 2.3 | 44.6 | 75.1 | 2.1 | 24.9 | 87.7 | 0.3 | 12.3 | 95.5 | 0.0 | 4.5 |
| U2-HiConf | 0.7 | 0.1 | 99.3 | 23.7 | 0.6 | 76.3 | 52.6 | 0.0 | 47.4 | 72.5 | 0.1 | 27.5 | 86.3 | 0.0 | 13.7 |
| UCHIME | 14.6 | 1.0 | 85.4 | 58.5 | 2.3 | 41.5 | 77.2 | 2.9 | 22.8 | 88.3 | 0.2 | 11.7 | 94.2 | 0.0 | 5.8 |
| ChSlayer | 2.8 | 0.0 | 97.2 | 29.7 | 0.8 | 70.3 | 54.4 | 1.5 | 45.6 | 68.9 | 2.0 | 31.1 | 87.8 | 0.7 | 12.2 |
| ChSlayer-kmer | 3.6 | 0.1 | 96.4 | 32.1 | 0.8 | 67.9 | 54.0 | 1.0 | 46.0 | 65.7 | 1.9 | 34.3 | 87.9 | 0.8 | 12.1 |
| ===== V5 ===== |  |  |  |  |  |  |  |  |  |  |  |  |  |  |  |
| Algo | Sens | S=90<br>FP | FN | Sens | S=95<br>FP | FN | Sens | S=97<br>FP | FN | Sens | S=99<br>FP | FN | Sens | S=100<br>FP | FN |
| U2-Denoised | 0.1 | 0.2 | 99.9 | 2.3 | 5.0 | 97.7 | 10.1 | 14.2 | 89.9 | 44.5 | 47.3 | 55.5 | 98.8 | 0.0 | 1.2 |
| U2-Sensitive | 69.4 | 20.0 | 30.6 | 82.4 | 40.6 | 17.6 | 85.0 | 25.6 | 15.0 | 88.4 | 6.3 | 11.6 | 93.7 | 0.0 | 6.3 |
| U2-Balanced | 31.2 | 2.7 | 68.8 | 63.1 | 8.6 | 36.9 | 71.9 | 3.1 | 28.1 | 79.7 | 0.2 | 20.3 | 84.7 | 0.0 | 15.3 |
| U2-Specific | 6.3 | 0.0 | 93.7 | 31.6 | 1.4 | 68.4 | 45.8 | 0.4 | 54.2 | 59.6 | 0.0 | 40.4 | 71.5 | 0.0 | 28.5 |
| U2-HiConf | 1.0 | 0.0 | 99.0 | 7.2 | 0.0 | 92.8 | 14.5 | 0.0 | 85.5 | 29.4 | 0.0 | 70.6 | 42.5 | 0.0 | 57.5 |
| UCHIME | 7.3 | 0.0 | 92.7 | 30.6 | 0.7 | 69.4 | 45.0 | 0.0 | 55.0 | 57.9 | 0.0 | 42.1 | 65.2 | 0.0 | 34.8 |
| ChSlayer | 0.8 | 0.0 | 99.2 | 5.8 | 0.2 | 94.2 | 14.3 | 0.6 | 85.7 | 24.5 | 0.0 | 75.5 | 28.2 | 0.1 | 71.8 |
| ChSlayer-kmer | 0.6 | 0.0 | 99.4 | 6.0 | 0.1 | 94.0 | 15.9 | 0.3 | 84.1 | 25.2 | 0.1 | 74.8 | 32.7 | 0.1 | 67.3 |
| ===== ITS1 ===== |  |  |  |  |  |  |  |  |  |  |  |  |  |  |  |
| Algo | Sens | S=90<br>FP | FN | Sens | S=95<br>FP | FN | Sens | S=97<br>FP | FN | Sens | S=99<br>FP | FN | Sens | S=100<br>FP | FN |
| U2-Denoised | 11.0 | 0.7 | 89.0 | 23.9 | 30.3 | 76.1 | 40.6 | 34.0 | 59.4 | 44.5 | 67.9 | 55.5 | 95.1 | 0.0 | 4.9 |
| U2-Sensitive | 7.9 | 3.7 | 92.1 | 29.1 | 9.2 | 70.9 | 46.7 | 7.1 | 53.3 | 64.6 | 2.6 | 35.4 | 82.7 | 0.0 | 17.3 |
| U2-Balanced | 26.6 | 10.9 | 73.4 | 53.3 | 15.6 | 46.7 | 66.9 | 10.6 | 33.1 | 78.2 | 3.1 | 21.8 | 92.7 | 0.0 | 7.3 |
| U2-Specific | 7.9 | 3.6 | 92.1 | 31.6 | 9.3 | 68.4 | 47.6 | 7.5 | 52.4 | 65.4 | 2.7 | 34.6 | 85.7 | 0.0 | 14.3 |
| U2-HiConf | 2.0 | 1.4 | 98.0 | 12.4 | 5.4 | 87.6 | 23.5 | 4.4 | 76.5 | 48.2 | 1.4 | 51.8 | 67.6 | 0.0 | 32.4 |
| UCHIME | 8.3 | 1.9 | 91.7 | 29.7 | 3.3 | 70.3 | 49.2 | 1.6 | 50.8 | 67.8 | 0.4 | 32.2 | 78.2 | 0.0 | 21.8 |
| ChSlayer | (ChimeraSlayer does not support ITS) |  |  |  |  |  |  |  |  |  |  |  |  |  |  |
| ChSlayer-kmer | (ChimeraSlayer does not support ITS) |  |  |  |  |  |  |  |  |  |  |  |  |  |  |
| ===== ITS2 ===== |  |  |  |  |  |  |  |  |  |  |  |  |  |  |  |
| Algo | Sens | S=90<br>FP | FN | Sens | S=95<br>FP | FN | Sens | S=97<br>FP | FN | Sens | S=99<br>FP | FN | Sens | S=100<br>FP | FN |
| U2-Denoised | 0.0 | 8.5 | 100.0 | 15.2 | 23.7 | 84.8 | 46.8 | 50.4 | 53.2 | 50.4 | 76.1 | 49.6 | 97.0 | 0.0 | 3.0 |
| U2-Sensitive | 3.9 | 1.0 | 96.1 | 22.7 | 2.5 | 77.3 | 43.0 | 2.4 | 57.0 | 65.4 | 0.3 | 34.6 | 76.7 | 0.0 | 23.3 |
| U2-Balanced | 17.2 | 5.4 | 82.8 | 50.4 | 8.0 | 49.6 | 67.8 | 6.1 | 32.2 | 82.3 | 0.9 | 17.7 | 90.1 | 0.0 | 9.9 |
| U2-Specific | 3.5 | 1.2 | 96.5 | 24.4 | 2.9 | 75.6 | 44.3 | 2.7 | 55.7 | 67.6 | 0.5 | 32.4 | 81.1 | 0.0 | 18.9 |
| U2-HiConf | 0.6 | 0.1 | 99.4 | 6.7 | 1.3 | 93.3 | 19.8 | 1.3 | 80.2 | 45.1 | 0.1 | 54.9 | 63.3 | 0.0 | 36.7 |
| UCHIME | 4.4 | 0.6 | 95.6 | 28.4 | 0.9 | 71.6 | 46.5 | 0.6 | 53.5 | 65.9 | 0.1 | 34.1 | 74.2 | 0.0 | 25.8 |
| ChSlayer | (ChimeraSlayer does not support ITS) |  |  |  |  |  |  |  |  |  |  |  |  |  |  |
| ChSlayer-kmer | (ChimeraSlayer does not support ITS) |  |  |  |  |  |  |  |  |  |  |  |  |  |  |

**Fig. SN12.1.**
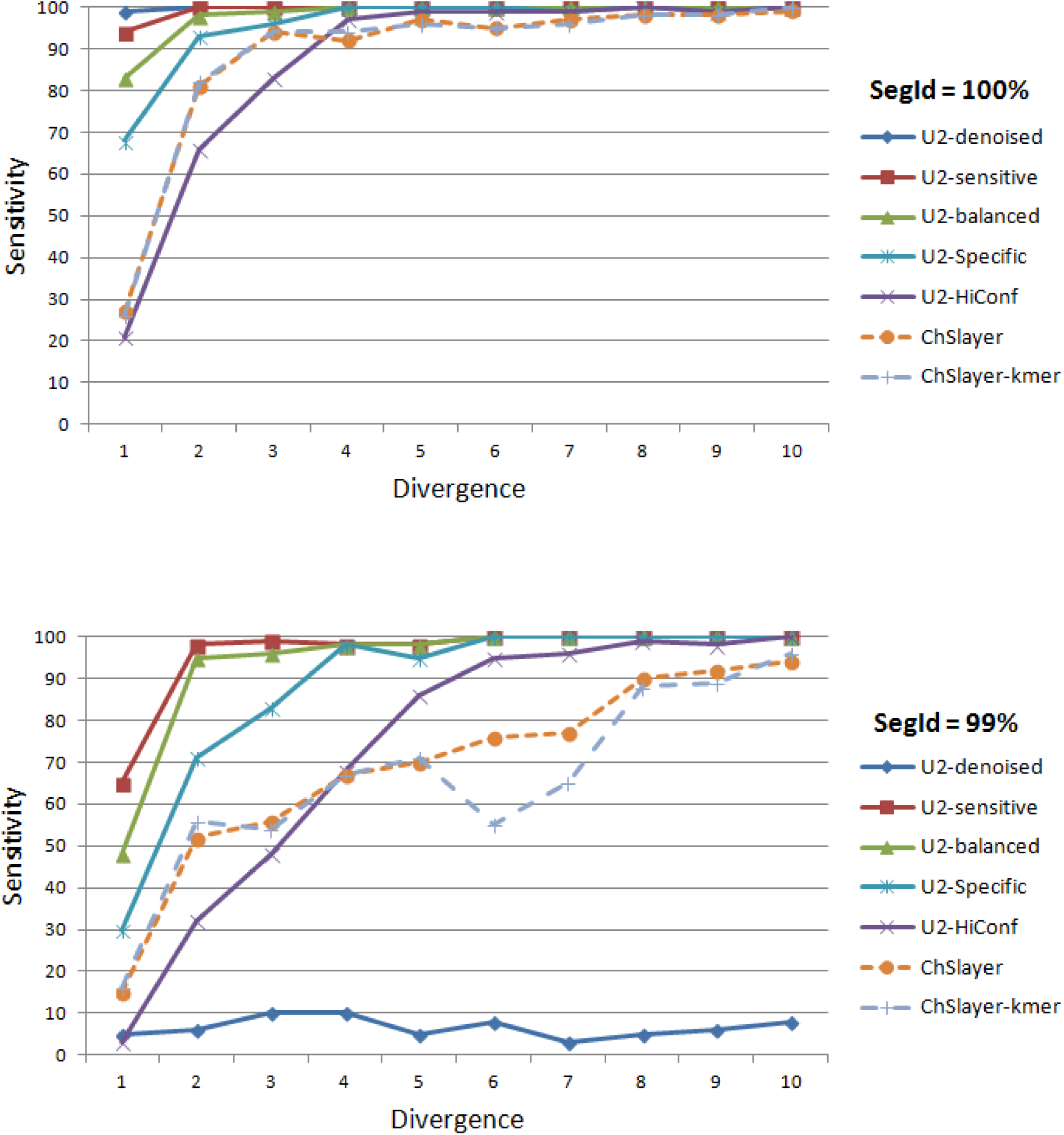
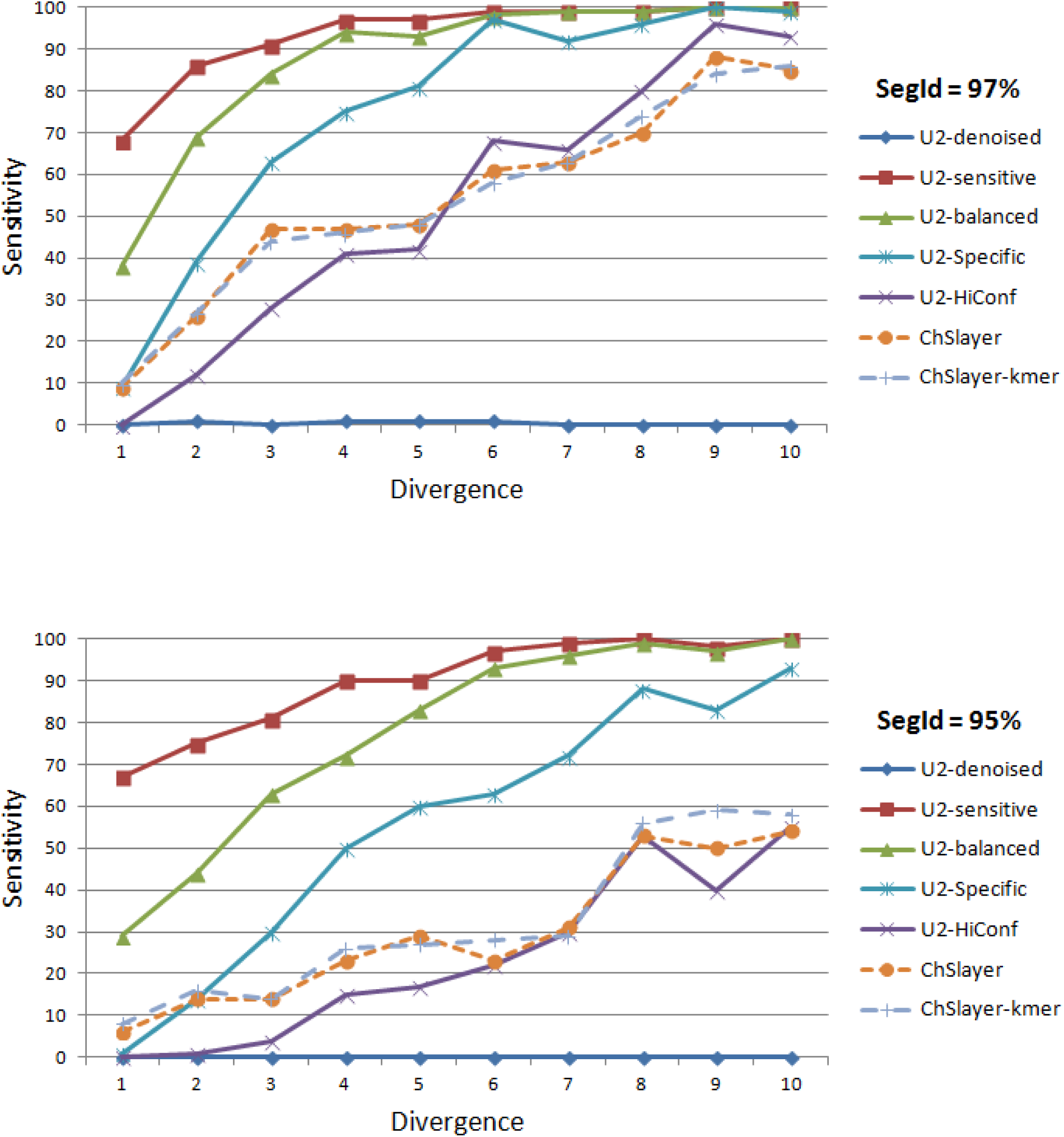
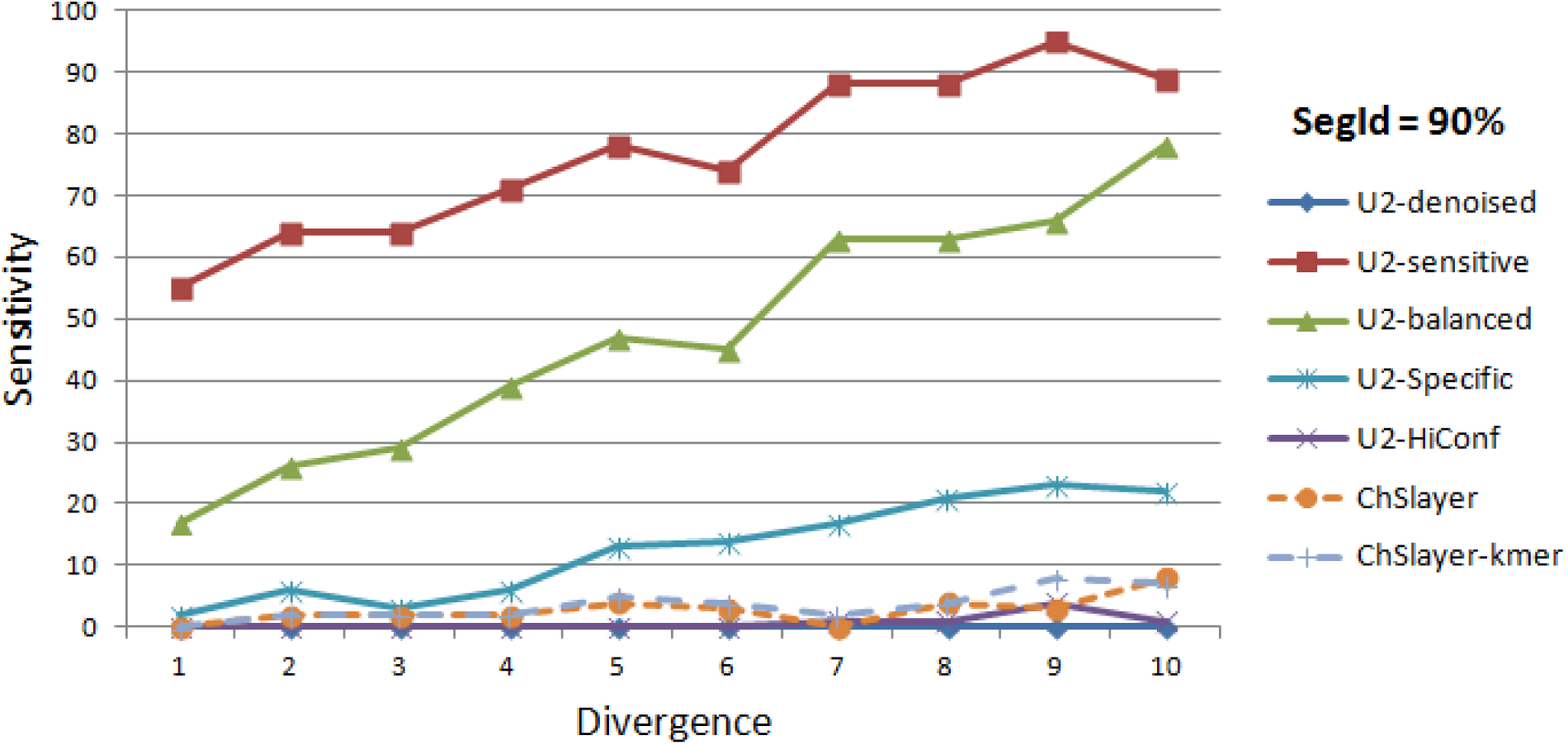
Variation of sensitivity on the V4 region with *D* and *S*. The charts show sensitivity as a function of divergence for the V4 region measured on CHSIMA with segment identity *S* = 100, 99, 97, 95 and 90% and divergence *D* = 1, 2 … 10. Sensitivities are given as percentages while divergences are substitutions, so e.g. *D* = 10 corresponds to a fractional divergence of (10 / |V4|) = 10/250 = 4%,

## References

1. HMP Consortium & Notes, S. A framework for human microbiome research. Nature 486, 215–21 (2012).

2. DeKosky, B. J. et al. High-throughput sequencing of the paired human immunoglobulin heavy and light chain repertoire. Nat. Biotechnol. 31, 166–9 (2013).

3. Gagan, J. & Van Allen, E. M. Next-generation sequencing to guide cancer therapy. Genome Med 7, 80 (2015).

4. Haas, B., Gevers, D. & Earl, A. Chimeric 16S rRNA sequence formation and detection in Sanger and 454-pyrosequenced PCR amplicons. Genome Res. 494–504 (2011). doi:10.1101/gr.112730.110.Freely

5. Edgar, R. C., Haas, B. J., Clemente, J. C., Quince, C. & Knight, R. UCHIME improves sensitivity and speed of chimera detection. Bioinformatics 27, 2194–200 (2011).

6. Wright, E. S., Yilmaz, L. S. & Noguera, D. R. DECIPHER, a search-based approach to chimera identification for 16S rRNA sequences. Appl. Environ. Microbiol. 78, 717–25 (2012).

7. Mysara, M., Saeys, Y., Leys, N., Raes, J. & Monsieurs, P. CATCh, an ensemble classifier for chimera detection in 16s rRNA sequencing studies. Appl. Environ. Microbiol. 81, 1573–1584 (2015).

8. Edgar, R. C. & Flyvbjerg, H. Error filtering, pair assembly and error correction for next-generation sequencing reads. Bioinformatics 31, 3476–3482 (2014).

9. Edgar, R. C. Search and clustering orders of magnitude faster than BLAST. Bioinformatics 26, 2460–1 (2010).

10. Pruesse, E. et al. SILVA: A comprehensive online resource for quality checked and aligned ribosomal RNA sequence data compatible with ARB. Nucleic Acids Res. 35, 7188–7196 (2007).

11. Tikhonov, M., Leach, R. W. & Wingreen, N. S. Interpreting 16S metagenomic data without clustering to achieve sub-OTU resolution. ISME J. 9, 68–80 (2015).

12. Callahan, B. J. et al. DADA2 : High resolution sample inference from amplicon data. bioRxiv 0–14 (2015). doi:10.1101/024034

13. Lahr, D. J. G. & Katz, L. a. Reducing the impact of PCR-mediated recombination in molecular evolution and environmental studies using a new-generation high-fidelity DNA polymerase. Biotechniques 47, 857–66 (2009).

14. Edgar, R. C. UPARSE: highly accurate OTU sequences from microbial amplicon reads. Nat. Methods 10, 996–8 (2013).

## Supplementary References

1. Kozich, J. J., Westcott, S. L., Baxter, N. T., Highlander, S. K. & Schloss, P. D. Development of a dual-index sequencing strategy and curation pipeline for analyzing amplicon sequence data on the miseq illumina sequencing platform. Appl. Environ. Microbiol. 79, 5112–5120 (2013).

2. Bokulich, N. A. et al. Quality-filtering vastly improves diversity estimates from Illumina amplicon sequencing. Nat. Methods 10, 57–9 (2013).

3. Edgar, R. C. & Flyvbjerg, H. Error filtering, pair assembly and error correction for next-generation sequencing reads. Bioinformatics 31, 3476–3482 (2014).

4. Callahan, B. J. et al. DADA2: High resolution sample inference from amplicon data. bioRxiv 0–14 (2015). doi:10.1101/024034

5. Mysara, M., Saeys, Y., Leys, N., Raes, J. & Monsieurs, P. CATCh, an ensemble classifier for chimera detection in 16s rRNA sequencing studies. Appl. Environ. Microbiol. 81, 1573–1584 (2015).

6. Edgar, R. C., Haas, B. J., Clemente, J. C., Quince, C. & Knight, R. UCHIME improves sensitivity and speed of chimera detection. Bioinformatics 27, 2194–200 (2011).

7. Wright, E. S., Yilmaz, L. S. & Noguera, D. R. DECIPHER, a search-based approach to chimera identification for 16S rRNA sequences. Appl. Environ. Microbiol. 78, 717–25 (2012).

8. Haas, B., Gevers, D. & Earl, A. Chimeric 16S rRNA sequence formation and detection in Sanger and 454-pyrosequenced PCR amplicons. Genome Res. 494–504 (2011). doi:10.1101/gr.112730.110.Freely

9. Pruesse, E. et al. SILVA: A comprehensive online resource for quality checked and aligned ribosomal RNA sequence data compatible with ARB. Nucleic Acids Res. 35, 7188–7196 (2007).

10. DeSantis, T. Z. et al. Greengenes, a chimera-checked 16S rRNA gene database and workbench compatible with ARB. Appl. Environ. Microbiol. 72, 5069–72 (2006).

11. Schloss, P. D. et al. Introducing mothur: open-source, platform-independent, community-supported software for describing and comparing microbial communities. Appl. Environ. Microbiol. 75, 7537–41 (2009).

